# Lipidome of Compact Myelin from Cortical Regions of the Aging Murine Brain

**DOI:** 10.64898/2026.11.01.763492

**Authors:** Victor P. Liu, Connor E. Johnson, Shaheer Lakhani, Kendra L. Furber

## Abstract

Age-related cognitive decline has been associated with neurodegeneration in cortical white matter tracts. The trajectory of white matter degeneration over the lifespan follows a similar pattern in humans, primates and rodents including the loss of myelin integrity and concomitant loss in myelinated axonal segments. Previous studies suggest that biochemical alterations in myelin begin to occur well prior to myelin loss. To better understand these biochemical determinants underlying white matter degeneration, we investigated the lipid composition of myelin isolated from young adult, middle-aged and aged murine brain. Compact myelin was enriched from cortical regions using a refined density centrifugation protocol and the lipidome was profiled using LC-MS/MS. Over 2000 lipid species were identified, with the relative expression of 414 species changing over the age-span. The myelin lipidome from young mice was biochemically distinct compared to the aging cohorts. Aging was characterized by an overall decrease in high abundant species of glycerophospholipid and sphingolipids, including phosphocholine, phosphoethanolamine, sphingomyelin and cerebrosides. Pathway analysis by LipidSig 2.0 and lipid ontology analysis by LION/web highlighted alterations to lipid metabolism and membrane biophysical properties related to bilayer stability and organization. These data support the concept that myelin lipid compositional changes that occur by middle age lead to structural changes in myelin at later ages. The findings provide greater insight into age-related biochemical alterations in the myelin sheath and may aid in strategies to mitigate white matter degeneration over the lifespan.

## INTRODUCTION

Myelin is a highly ordered, compact multi-lamellar membrane that acts as insulator for neuronal axons enabling the fast, efficient transmission of nerve impulses. In neurodevelopment, myelination begins caudally in the spinal cord and hindbrain during embryonic stages and in forebrain regions during the postnatal period (1). Myelination continues into adolescence and adulthood, particularly in the cortical areas, contributing to plasticity that underlies learning and memory (2, 3). Over the human life span, volume of white matter in the brain peaks in middle-age and then starts to decline (4). This includes both a decrease in myelin integrity and a loss of myelin internodes in cortical regions, which has been correlated to age-related cognitive decline (5–7).

Age-related white matter degeneration has been shown to follow similar trajectories in rodent models (8). Oligodendrocytes, the myelinating glia of the CNS, are thought to become susceptible to cellular stress with age due to the high metabolic demand for maintaining the myelin sheath (9). Oligodendrocytes produce large amounts of highly organized membrane, myelinating up to 40-50 axons in cortical regions with a total membrane surface of up to 2mm^2^ / cell (10). Myelin is thought to be prone to age-related biochemical modifications, owing in part to the high lipid content of the myelin membranes and slow lipid turnover rate (11).

Ultrastructural analyses have shown a breakdown in myelin structure and a concomitant loss in myelinated axonal segments in aged mice (8, 12). Moreover, Fourier-transform infrared (FTIR) spectroscopy imaging of the corpus callosum in the aging mouse brain showed alterations in lipid content that was evident by middle-age prior to myelin structural changes (8).

Regional differences in glycerophospholipids, sphingolipids and sterols have been reported in the cortex and corpus callosum from the aging mouse brain (13–15). Oligodendrocytes show enrichment in glycolipids compared to other types of brain cells (14), and lipidomic analyses of myelin has previous identified over 600 lipid species (16). Early biochemical studies on myelin membranes indicate alterations in lipid content across total phospholipids, sphingolipids and cholesterol over the age-span (17–20); however, the advancement in mass spectrometric techniques has the potential to characterize numerous different species within each subclass.

Here, we have employed a global lipidomic platform to identify alterations in lipid composition in isolated myelin membranes from murine brain cortex, including the underlying white matter of the corpus callosum, from across the lifespan. Age-related changes in myelin lipid composition, along with myelin structural proteins, are discussed in the context of membrane structure and function.

## RESULTS

### Characterization of isolated myelin fractions

To investigate biochemical alterations that underly myelin degeneration in the aging brain, myelin membranes were isolated using a well-established method of density centrifugation (Fig 1A). Myelin fractions from whole brain were initially assessed for the enrichment of canonical myelin markers (Fig 1B). The major myelin proteins (21), such as myelin proteolipid protein (PLP), myelin basic protein (MBP), 2′, 3′-cyclic nucleotide 3′-phosphodiesterase (CNP), sirtuin 2 (SIRT2) and oligodendrocyte glycoprotein (MOG), were enriched in the crude compact myelin fraction from the first gradient, as well as refined myelin fractions from the second gradient. On the other hand, myelin proteins associated with paranodal regions, such as neurofascin 155 (NF155), were enriched in the crude compact myelin fraction from the first gradient but showed greater enrichment in the higher density myelin fraction from the second gradient. Next the myelin fractions were assessed for purity through exclusionary markers specific for other CNS cell types. The astrocytic glial fibrillary acid protein (GFAP), the microglial ionized calcium-binding adaptor molecule (IBA1) and the axonal proteins neurofilament heavy (NEFH) and spectrin beta (SPTBN1) were found in crude compact myelin but then subsequently fractionated out of the myelin light and medium in the second gradient. The cytoskeletal protein tubulin beta-3 (TUBB3) and tubulin beta-4 (TUBB4) were present across most fractions, but to a much lesser degree in compact myelin light and medium. The exclusionary markers for other cellular compartments, such as endoplasmic reticulum (GRP78), mitochondria (ATP5a) and nodes of Ranvier (Na^+^/K^+^ ATPase), were also diminished in myelin light and medium fractionations in the second gradient. Thus, the 2-gradient fractionation method shows reliable enrichment of myelin membranes.

**Figure 1.**
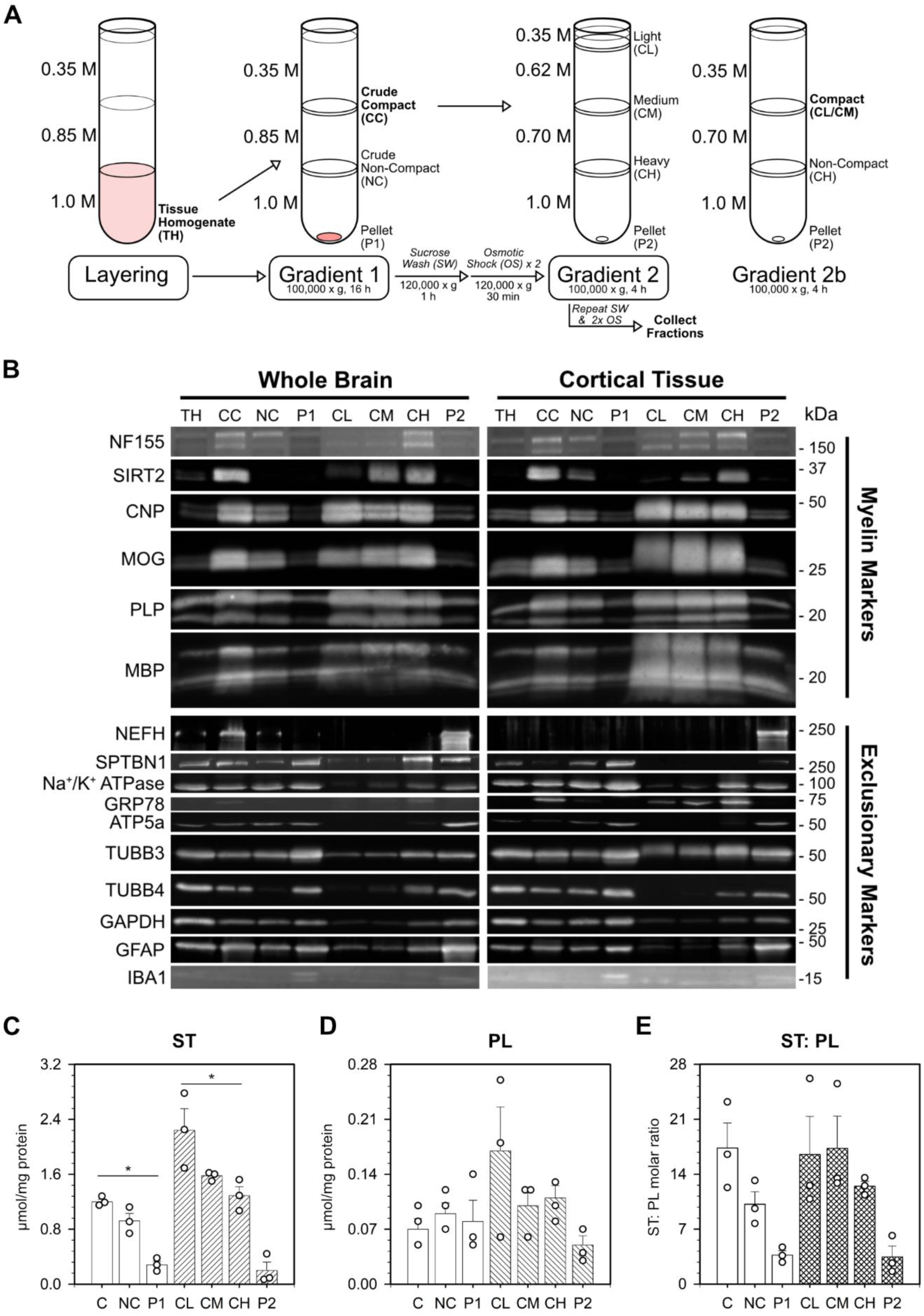
Enrichment of compact myelin from the murine brain. (**A**) Schematic of myelin isolation protocol using a discontinuous sucrose gradient. Mouse brain homogenate (TH) was separated on gradient 1 to produce crude compact myelin (CC; 0.35 M/0.85 M interface), non-compact myelin (NC; 0.85M/1.0M interface) and pellet (P1). Gradient 2 was used to separate the crude compact myelin into light (CL; 0.32M/0.62M interface), medium (CM; 0.62M/0.70M interface) and heavy (CH; 0.70M/1.0M interface) fractions and pellet (P2). A modified second gradient (2b) was used to combine CL / CM into single compact myelin fraction. (**B**) Immunoblotting was performed on myelin fractions enriched from whole brain or cortical tissue homogenate for myelin markers (NF155, SIRT2, CNP, MOG, MBP and PLP) and exclusionary markers (NEFH, SPTBN1, Na^+^/K^+^-ATPase, GRP78, ATP5A, TUBB3, TUBB4, GAPDH, GFAP and IBA1) to assess isolation method. (**C-E**) Biochemical assays for sterol (C; ST), phospholipid (D; PL) and ST: PL ratio (E) showed enrichment of myelin subfractions. **Statistical analyses**: Data represented by biological replicates (n = 3), with mean ± SEM; Kruskal-Wallis test (*p < 0.05).

As the age-related loss of myelin is has been primarily observed in cortical white matter tracts (5–7), the fractionation method was also validated with cortical tissue that included the cortex and underlying corpus callosum. Western blot analysis of the myelin proteins and exclusionary markers from myelin isolated from cortical tissue showed similar pattern to that of myelin isolated from whole brain (Fig 1B). Along with myelin structural proteins, myelin membranes are enriched in lipid content particularly in cholesterol (18, 20). When looking at the lipid distribution, sterol (ST; Fig 1C), phospholipid (PL; Fig 1D) and the ST:PL ratio (Fig 1E) increased across all the myelin fractions. Based on the similar distribution of protein and lipid composition of compact myelin light and compact myelin medium, these were combined into a single compact myelin fraction on a modified second gradient (Fig 1A; gradient 2b CL/CM) to generate ample material for downstream global lipidome analysis.

### Lipidome of the compact myelin from the aging brain

The lipid composition of myelin will contribute to the structural integrity of this highly organized membrane (22). To further explore this, untargeted lipidomics was performed on compact myelin isolated from the cortical regions of male and female mice at different timepoints across the lifespan: young, middle-age and aged (n = 4; n = 2 per sex). The normalized intensity for peaks with the same molecular identification were combined to simplify the biological interpretation of the data, which resulted in identification for a total of 2050 lipid species across the different subclasses (Fig 2A; Table S1; 601 Tier 1 IDs, 161 Tier 2 IDs, and 1288 Tier 3 IDs). Dimensional reduction by PCA showed distinct separation of the myelin lipidome across age groups (Fig 2B). Further analyses revealed only marginal differences in sample clustering patterns based on sex or with exclusion of Tier 3 identifications (Fig S1). Volcano plots of the relative expression of lipid species across age groups indicate a greater number of changes between middle-age vs. young (Fig 2C; 136 down, 178 up) and aged vs. young (Fig 2D; 165 down, 225 up) compared to aged vs. middle-age (Fig 2E; 21 down, 45 up). The most altered lipid classes were the main components of biological membranes, including glycerophospholipids and sphingolipids (Fig 2F). To better understand the contribution of individual lipids, the topmost abundant species from each class were further examined.

**Figure 2.**
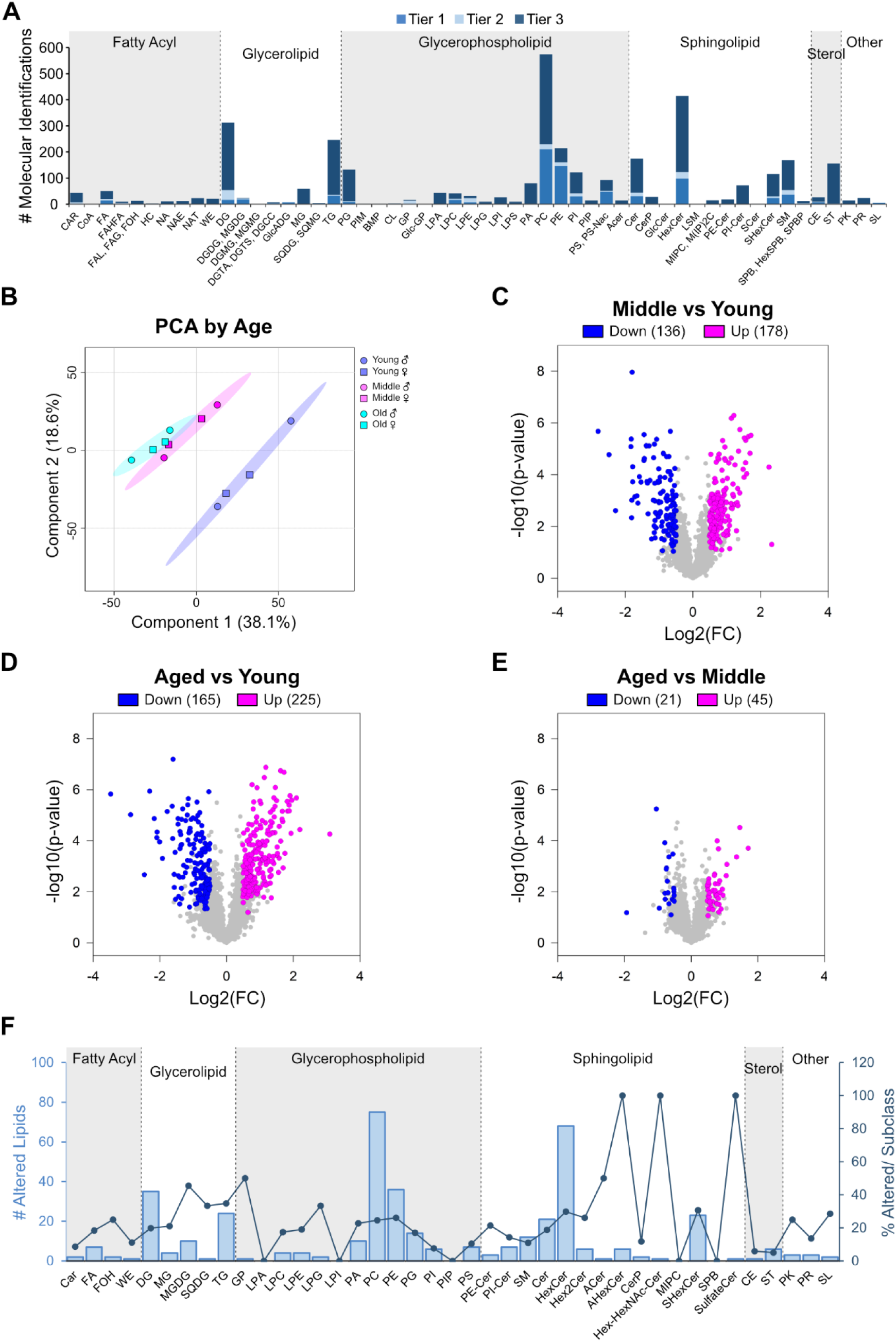
Global characterization of lipidome in murine cortical myelin over the age-span. (**A**) Number of unique molecular identifications from MS/MS spectra putatively identified across different lipid classes and subclasses (n = 4; n = 2/sex). Features identified as the same molecular lipid species were combined for multivariate and univariate analysis. (**B**) PCA score plot displaying age-dependent clustering of unique molecular identifications in myelin lipidome from young (3 months; blue), middle-aged (12 months; magenta), and aged (20 months; cyan). (**C-E**) Volcano plots depict the number of altered lipid species for middle-age vs young (C), aged vs young (D) and aged vs middle-aged mice (E). (**F**) Representation of the number and percentage of altered lipid species across different lipid subclasses. **Statistical analyses**: one-way ANOVA followed by Fisher LSD post-hoc test. Comparisons were considered significant at p < 0.05 after false discovery rate (FDR) correction (q < 0.1).

Glycerophospholipids are the backbone of biological membranes forming the majority of the lipid bilayer (23). The percentage of identified species that showed changes in abundance between age groups for phosphatidylcholine (PC), phosphatidylethanolamine (PE), phosphatidic acid (PA), phosphatidylglycerol (PG), phosphatidylserine (PS), and phosphatidylinositol (PI/PIP) were 25%, 26%, 23%, 17%, 10%, and 7%, respectively (Fig 2F). The relative abundance of glycerophospholipid subclasses over the age span suggests minimal changes in the overall distribution of headgroups, except for PC which decreased by ∼6% (Fig 3A). The most abundant lipid species with altered expression levels for PC (Fig 3B) and ether-linked PC (Fig 3C) all decreased over the age span. Similarly, the majority of most abundant lipid species with altered expression levels for PE (Fig 3D) and ether-linked PE (Fig 3E) also decreased. Other phospholipid classes showed a variety of species that decreased and increased in abundance (Fig 3F).

**Figure 3.**
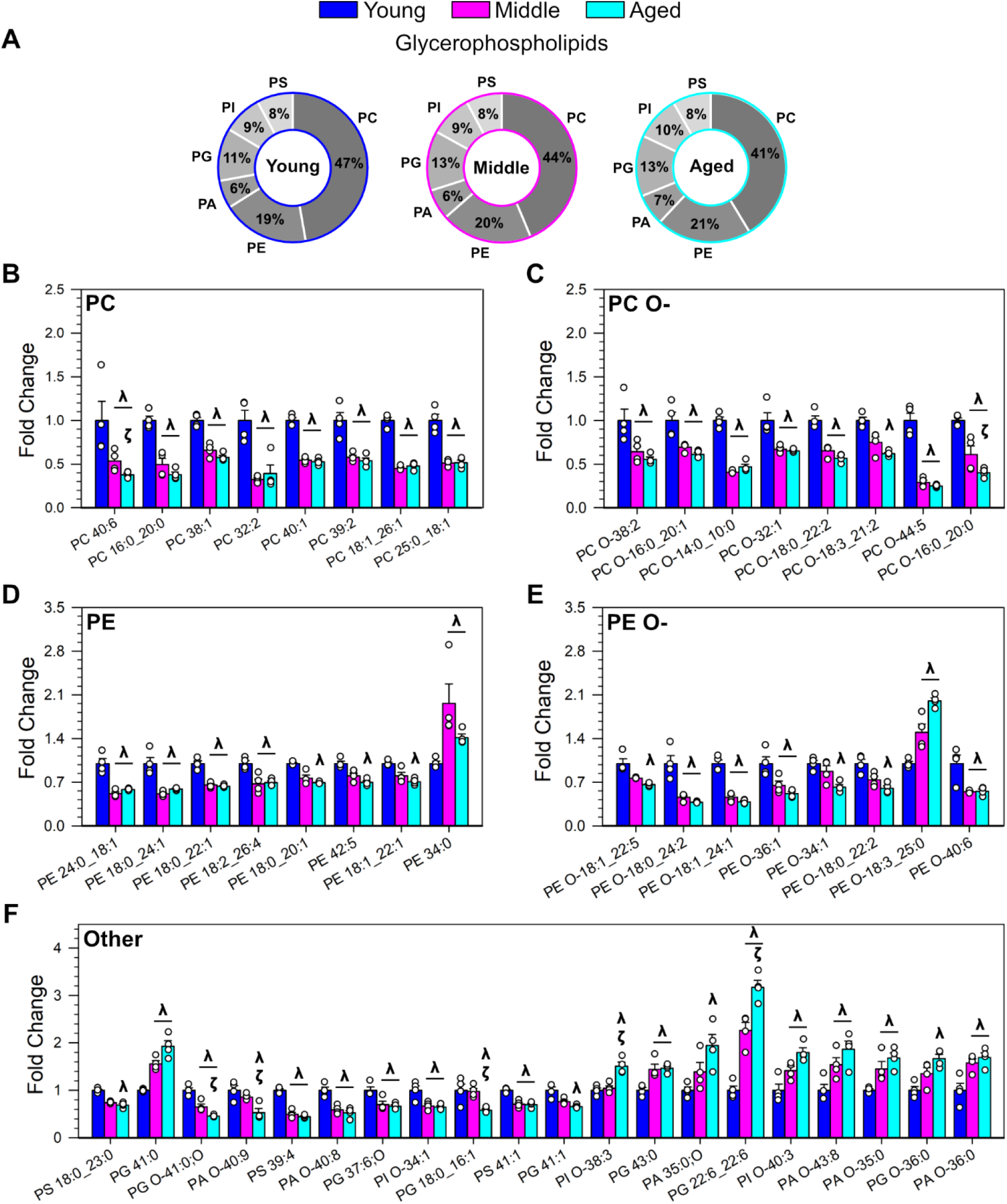
Age-dependant alterations in glycerophospholipids in murine cortical myelin. (**A**) Relative abundance (%) of glycerophospholipids in the myelin lipidome from young (3 months; blue), middle-aged (12 months; magenta) and aged (20 months; cyan) mice. (**B-F**) The top 8 most abundant lipid species that showed age-dependent changes in expression for each glycerophospholipid subclass are depicted: phosphatidylcholine (B; PC), ether-linked phosphatidylcholine (C; PC O-), phosphatidylethanolamine (D; PE), and ether-linked phosphatidylethanolamine (E; PE O-). The top 20 most abundant glycerophospholipids in other classes: phosphatidyl serine (PS), phosphatidylglycerol (PG), ether-linked PG (PG O-), phosphatidic acid (PA), ether-linked PA (PA O-), and ether-linked phosphatidylinositol (PI O-). **Statistical analyses**: Normalized intensities expressed as fold change relative to young age group. Data represented as biological replicates (n = 4; n = 2/sex), with mean ± SEM; one-way ANOVA followed by Fisher LSD post-hoc test. Comparisons were considered significant (**^λ^**p < 0.05 vs young, **^ζ^**p < 0.05 vs middle-aged) after false discovery rate (FDR) correction (q < 0.1).

Sphingolipids that contain a ceramide backbone are enriched in myelin membranes (22). Sphingomyelin (SM), along with glycerophospholipids, make up the lipid bilayer while preferentially interacting with sterols (ST), like cholesterol, to form lipid domains (23). The relative abundance of sphingolipid subclasses over the age span suggests a modest decrease in SM (Fig 4a), along with a shift from ST to CE (Fig 5A). For SM, 11% of the identified species showed changes in abundance between age groups whereas only 4% of ST species changed (including sterol esters or cholesterol esters, CE) (Fig 2G). Other subclasses of sphingolipid showed a greater percentage of identified species that were altered including 19% of ceramides (Cer), 30% of cerebrosides (hexosylceramides; HexCer) and 31% of sulfatides (sulfohexosylceramides; SHexCer) (Fig 2G). The most abundant species with altered expression levels over the age span showed a variety of changes, although there was an overall trend in sphingolipids being decreased (Fig 4B-E) and sterols being increased (Fig 5B).

**Figure 4.**
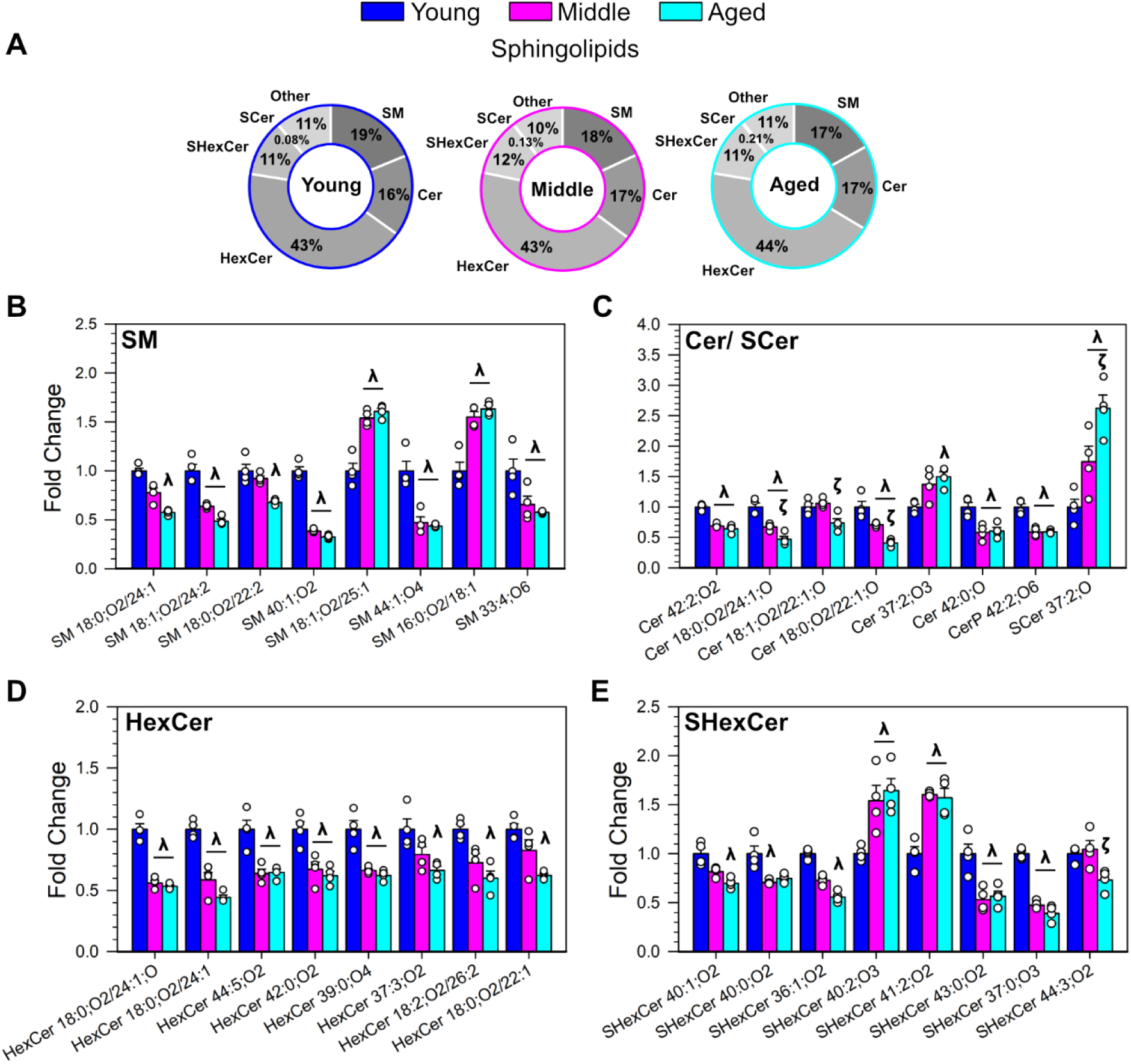
Age-dependant alterations in sphingolipids in murine cortical myelin. (**A**) Relative abundance (%) of sphingolipids in the myelin lipidome from young (3 months; blue), middle-age (12 months; magenta) and aged (20 months; cyan) mice. (**B-E**) The top 8 most abundant lipid species that showed age-dependent changes in expression for each sphingolipid subclass are depicted: sphingomyelin (B; SM), ceramides and sulfoceramides (C; Cer, SCer), hexosylceramides (D; HexCer), and sulfohexosylceramides (E; SHexCer). **Statistical analyses**: Normalized intensities expressed as fold change relative to young age group. Data represented as biological replicates (n = 4; n = 2/sex), with mean ± SEM; one-way ANOVA followed by Fisher LSD post-hoc test. Comparisons were considered significant (**^λ^**p < 0.05 vs young, **^ζ^**p < 0.05 vs middle-aged) after false discovery rate (FDR) correction (q < 0.1).

**Figure 5.**
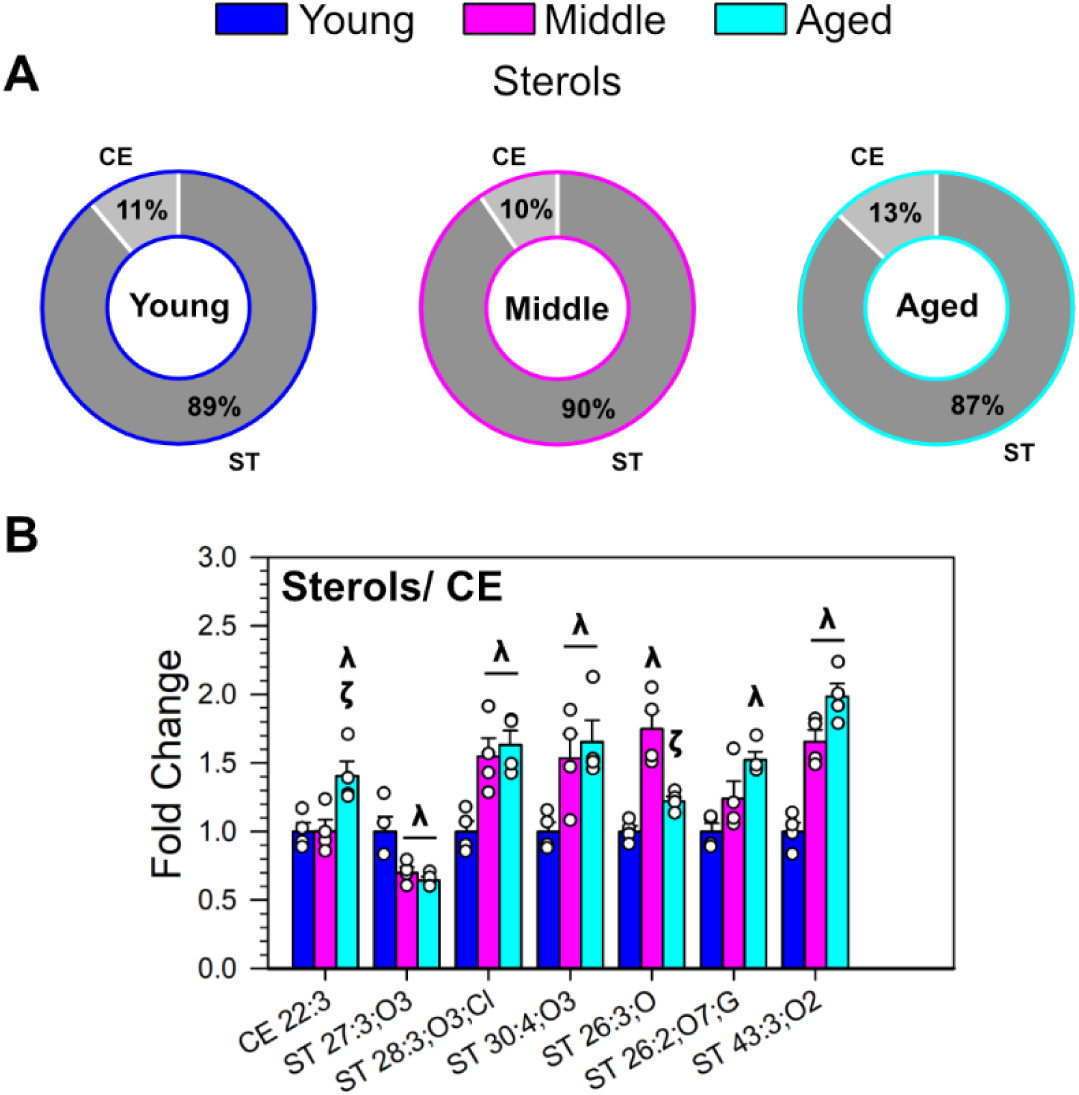
Age-dependant alterations in sterols in murine cortical myelin. (**A**) Relative abundance (%) of sterols in the myelin lipidome from in young (3 months; blue), middle-age (12 months; magenta) and aged (20 months; cyan) mice. (**B**) The top 7 most abundant lipid species that showed age-dependent changes in expression for sterol (ST) and cholesterol esters (CE) are depicted. **Statistical analyses**: Normalized intensities expressed as fold change relative to young age group. Data represented as biological replicates (n = 4; n = 2/sex), with mean ± SEM; one-way ANOVA followed by Fisher LSD post-hoc test. Comparisons were considered significant (**^λ^**p < 0.05 vs young, **^ζ^**p < 0.05 vs middle-aged) after false discovery rate (FDR) correction (q < 0.1).

### Effects of aging on myelin lipid composition, structure and function

To explore potential alterations in lipid metabolism that may influence the myelin lipidome during aging, LipidSig 2.0 was used to investigate lipid metabolic pathways in different glycerophospholipid, sphingolipid and glycerolipid subclasses (Fig 6a; Table S2). Overall, there was upregulation in conversion of phospholipids → lysophospholipids. There was also upregulation in conversion of PC → PS, PC → PA, and PC → SM with corresponding suppression of the reverse pathways across aging. For sphingolipid pathways, there was a shift in the balance between ceramides (Cer) and ceramide phosphates (CerP), as well as increase in conversion of cerbrosides (HexCer) to more complex glycolipids (Hex2Cer). Furthermore, there was a consistent trend across lifespan for the conversion of glycerophospholipids (PC, PI) into glycolipids (diacylglycerol; DG).

**Figure 6.**
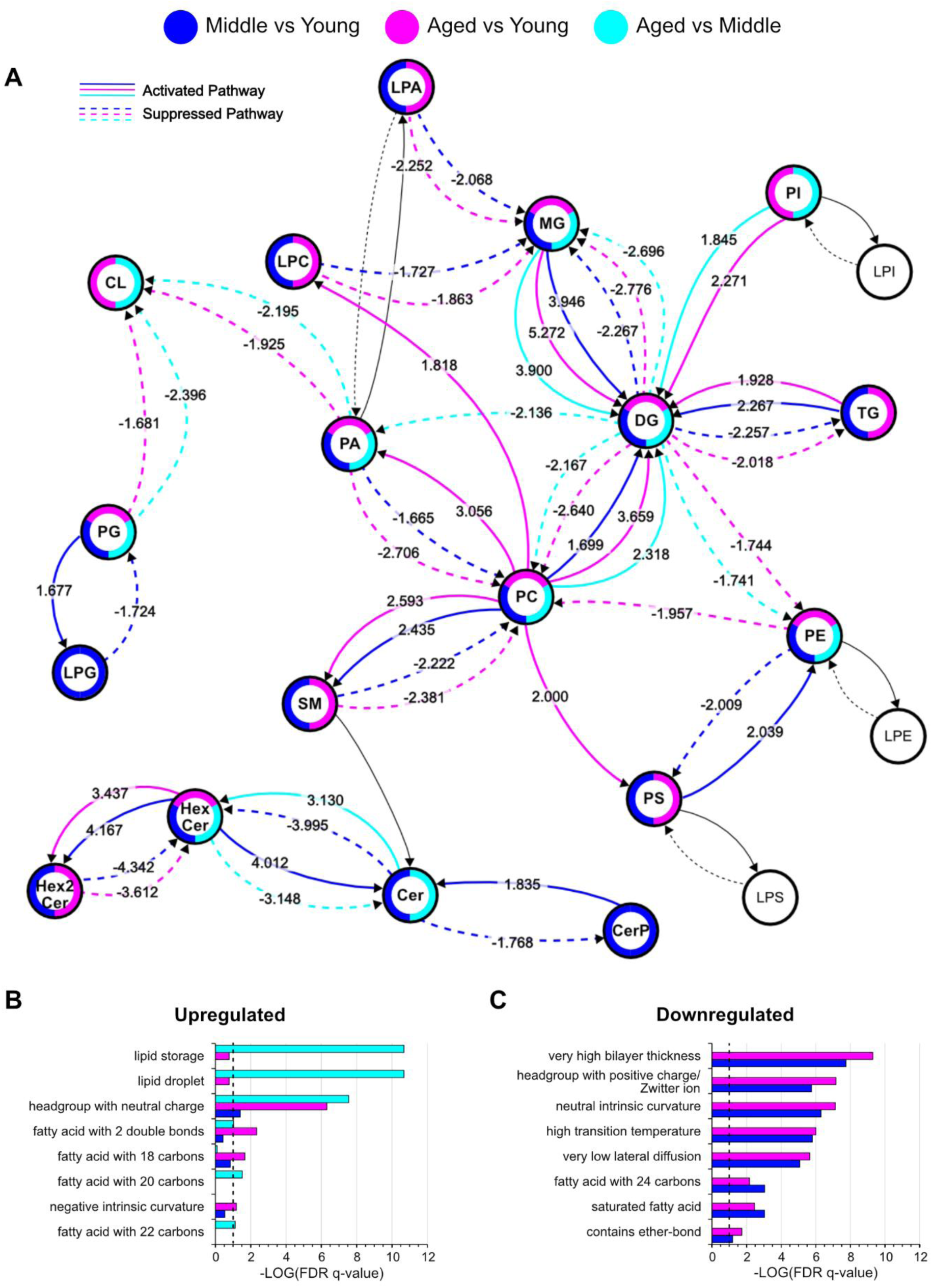
Predicted alterations in lipid pathways in murine cortical myelin lipidome over the age-span. (**A**) LipidSig 2.0 reaction network displaying metabolic alterations in synthesis pathways for middle-aged vs young (blue), aged vs young (magenta) and aged vs middle-aged (cyan) mice. Numbers denote the Z-score of activated (Z > 1.645, p-value < 0.05) and suppressed (Z < -1.645, p-value < 0.05) pathways shown in solid and dashed bold-colored lines, respectively. (**B-C**) Lipid ontology (LION) from LION/web show upregulated (B) and downregulated (C) terms related to membrane biophysical properties and lipid trafficking (FDR q < 0.1).

To understand the impact of lipidome alterations on the structure and function of myelin membranes, lipid ontology was performed using LION/web. The ontology terms could be broadly categorized into biochemical characteristics of lipids, biophysical characteristics of membranes, and lipid trafficking (Fig 6B-C; Fig S2; all terms including those that refer to specific lipid species or subclasses are reported in Table S3). For terms related to biochemical characteristics of lipids, the increase in headgroup with neutral charge with a corresponding decrease in headgroups positive / zwitterionic charge most likely reflects the decrease in the abundance of major PC and PE species. There was also a shift in fatty acid chain lengths from those most common for membrane lipids (18-22 carbons downregulated) to longer fatty acid chains (24 carbons upregulated) and a shift from saturated (downregulated) to more saturated fatty acids over lifespan, both of which would also affect membrane fluidity. For terms related to biophysical characteristics of membranes, there were alterations in bilayer thickness, decrease in transition temperature, increase in lateral diffusion and increase in intrinsic curvature during aging. Overall, these alterations in biophysical properties in the myelin membranes that may be indicative of altered myelin stability. Furthermore, there was an increase in lipid storage pathways observed in aged mice suggestive of altered lipid metabolism and transport.

### Impact on Myelin protein expression in the aging mouse brain

As membrane lipid composition may impact protein localization and function (24), the expression of the most abundant myelin proteins was assessed in myelin fractions from young middle-aged and aged mice (n = 4; n = 2 per sex). The overall recovery of compact and non-compact myelin fractions (by mass) did not change over aging (Fig 7A) There was no change in the expression of PLP (∼30 kDa; Fig 7B), an integral membrane protein which stabilizes the intraperiod line, or its isoform DM20 (∼26 kDa; not shown on graph). MBP, which stabilizes the major dense line, showed an increase of the predominate isoform (∼18.5 kDa; Fig 7C) in medium and heavy fractions, but there was not change in expression of other isoforms (∼21.5 kDa; not shown on graph). MOG, which resides on the outer lamellae of the sheath, increased in the heavy fraction only (Fig 7D). While there was no change in the expression of the phosphodiesterase CNP (Fig 7E), there was a large decrease in the deacetylase SIRT2 (Fig 7F). SIRT2 decreased in both the light and medium fractions from middle-age and aged mice. Thus, only modest changes in the expression of major structural proteins during aging, particularly in the compact myelin.

**Figure 7.**
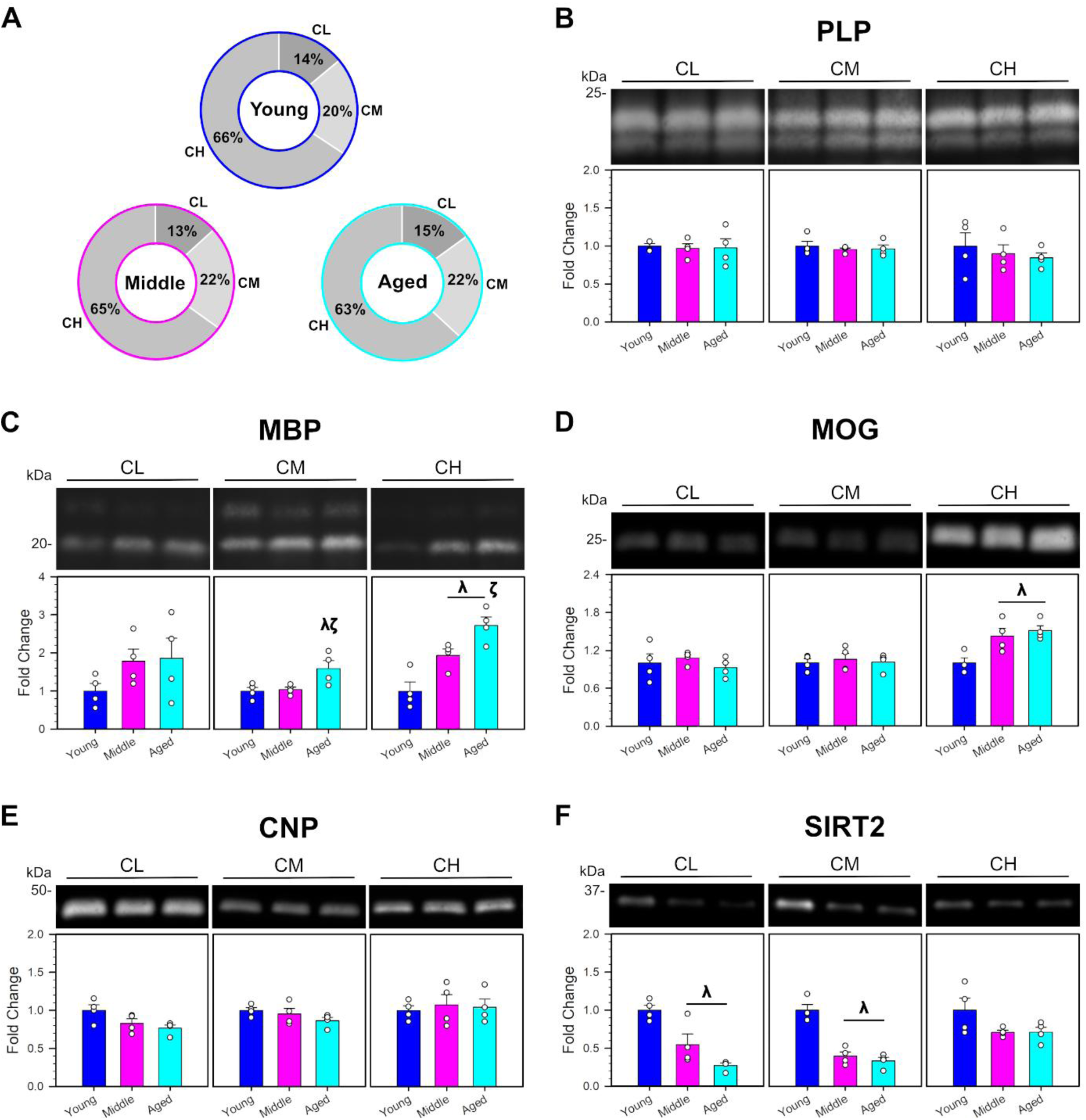
Expression of myelin proteins in murine cortical myelin over the age-span. **(A)** Mass distribution of isolated myelin subfractions was not altered in young (3 months), middle-aged (12 months) and aged (20 months) mice. (**B-F**) Immunoblot analysis of myelin protein expression during aging for PLP (B), MBP (C), MOG (D), CNP (E) and SIRT2 (F). **Statistical analyses**: Band volume was normalized to total protein and expressed as fold change relative to young age group. Data represented by biological replicates (n = 4; n = 2/sex), with mean ± SEM; one way ANOVA followed by Holm-Šidák post-hoc test (**^λ^**p < 0.05 vs young, **^ζ^**p < 0.05 vs middle-aged).

## 4. DISCUSSION

Myelin is continually remodelled over the lifespan based on experience (2, 3). The peak of white matter volume at middle-age (8) denotes a shift in the balance of de novo myelination and myelin degeneration. This is likely the consequence of a combination of metabolic changes in oligodendrocytes, mechanisms of membrane trafficking and turnover, and damage to phospholipids and proteins leading in changes to myelin composition. In this study, lipidomic analyses identified altered expression of numerous phospholipid, sphingolipid and sterol species that are the structural determinants in the formation of lipid bilayers. The altered lipid composition observed in myelin during aging impacts physicochemical properties of the membrane, such as thickness, fluidity and curvature, which also may directly or indirectly regulate protein function.

Heterogeneity in lipid composition between different biological membranes will determine their structure and function, particularly for specialized membranes such as the myelin sheath. Phospholipids, along with cholesterol, make up the backbone of the lipid bilayer (23). PC and SM, the most abundant glycerophospholipid and phosphosphingolipid in the brain (14), are associated with neutral curvature (25). The comparable size of the choline head group and fatty acid tails results in a cylindrical shape which leads to the formation of a flat bilayer. On the other hand, cholesterol is associated with negative curvature (26), where the small hydroxyl group and bulky hydrophobic ring structure form a conical shape. While adequate cholesterol is essential for proper myelination (27), high localized amounts of cholesterol may lead to regions of membrane deformation and instability (28). During aging, there was an increase in curvature of the myelin membrane which is likely reflective of the overall decrease in phospholipids and increase in sterols (and increase in glycerolipids, as discussed below).

Within individual biological membranes, there can also be considerable lateral heterogeneity through the formation of nanodomains (24). Sphingolipids promote tight packing of lipid bilayer due to their long, saturated fatty acid chains (23). They interact with cholesterol through dispersion forces and hydrogen bonding of the phosphate group to cholesterol hydroxyl group (29), which favors tight packing, increased order and decreased fluidity (23, 24). This arrangement also increases the thickness of the bilayer (24, 30). Oligodendrocytes (14) and their myelin membranes (18) are enriched in glycosphingolipids in the form of cerebrosides (HexCer) and sulfatides (SHexCer). Their long, saturated fatty acids promote domain organization while the sugar residues from apposing headgroups promote adhesion of the intraperiod line (31). Overall, there was a decrease in bilayer thickness, decrease in transition temperature and increase in lateral diffusion over the age-span that would all suggest less ordered myelin membrane.

The ordered, compact structure of myelin depends on the highly regulated interplay between membrane lipids and proteins. The formation of sphingolipid- and cholesterol-enriched membrane domains serve as a platform for protein-lipid and protein-protein interactions (24). Lipids may affect protein function directly by interaction with lipid binding domains or indirectly through the local microenvironment, such as altered conformation due the lateral pressure exerted by packing of acyl chains or altered orientation due to a mismatch between bilayer thickness and size of transmembrane domain (24). While expression levels of myelin proteins were not drastically changed with age, the altered lipid composition may affect the localization and functional role of these proteins. MBP and MOG were increased in non-compact myelin fractions, whereas SIRT2 was decreased in compact fractions. The trafficking of SIRT2 to the myelin sheath is dependent on PLP (32), an integral membrane protein that preferentially localizes to domains through its interaction with cholesterol (33). The addition of PLP into model membranes leads in rearrangement of lipid domains and aggregation of opposing bilayers (34), akin to its role in adhesion of the extracellular interface of myelin lamellae at the intraperiod line. At the major dense line, MBP facilitates the tight zippering of the cytoplasmic interface through strong electrostatic interactions with the negatively charged PS and PI (35–37). Thus, the organization of functional domains and distribution of charged lipids would be required for optimal compaction.

Unexpectedly, there were a number of glycerolipid species identified in isolated myelin which are not conventionally thought of as membrane lipids. The increase in glycerolipids, particularly DG, over the lifespans may have multiple impacts on cellular homeostasis. The abundance of DG in the membrane would be tightly regulated, both as a signaling molecule (38) and as its shape associated with high negative curvature (25). Furthermore, DG is also an important intermediate in triacylglycerol (TG) metabolism involved in lipid transport and storage (38). Pathways associated with lipid droplets were upregulated, largely due to alterations in TG and cholesterol (notably, with a shift in the relative abundance from sterols to sterol esters). An accumulation of lipid droplets has been observed in the aging mouse brain (39); however, it is uncertain whether these lipid droplets co-enrich with myelin membranes during the isolation process (40) or the possibility that glycerolipids accumulate within cytoplasmic pockets (23) in degenerating myelin. Regardless, these alterations are suggestive of aberrant lipid metabolism, and perhaps accumulation of lipids from degenerating myelin, that become apparent in later stages of life.

Alterations in the myelin lipidome were observed by middle-aged and persisted throughout the lifespan. A preliminary analysis of sex-differences showed only modest number of lipid species with altered abundance in myelin between male and female, yet with a limited sample size, more directed analysis of select lipid species would be warranted in future studies. The overarching trend was a decrease in phospholipids and sphingolipids, which are important structural components of the myelin membrane. This would lead to a loss of ordered arrangement of the lipid bilayer, decreased compaction of the lamellae and instability of the myelin sheath. The advances in mass spectrometry have highlighted the vast chemical diversity of lipids which are composed of various combinations of headgroups, backbone and fatty acid moieties. However, the biochemical analyses on isolated myelin membranes only give insight into “bulk” membrane composition. Limitations in methodology for studying lipid dynamics at the molecular level, particularly in biological membranes, do not address the intricacies of bilayer asymmetry, lateral heterogeneity and transient interactions within the lipid-protein mosaic. Regardless, the alterations in the myelin membrane composition identified here shed new light on the molecular underpinnings of age-dependent myelin degeneration. Importantly, these biochemical changes are apparent by middle-age prior to the loss of myelin indicating that early interventions are key to mitigating white matter degeneration during aging.

## MATERIALS AND METHODS

### Chemicals and Reagents

Reagents used for myelin membrane fractionation and antibodies used as inclusionary and exclusionary markers for validation are listed in Table 1 and 2, respectively.

**Table 1.** List of Basic Chemicals Used.

| **Chemical** | **Producer** | **Product Number** |
|---|---|---|
| 30% Acrylamide: Bis-acrylamide (37.5:1) | Bio-Rad | 1610158 |
| 4-(2-hydroxyethyl)-1-piperazineethanesulfonic acid (HEPES) | Fisher Scientific | BP310 |
| Acetic Acid | Fisher Scientific | A38 |
| Ammonium Persulfate | Fisher Scientific | BP179 |
| Ammonium Sulfate | Thermofisher Scientific | A11682.0B |
| Beta-mercaptoethanol | Bio-Rad | 1610710 |
| Bovine Serum Albumin | Sigma Aldrich | A9647 |
| Bromophenol Blue | Thermofisher Scientific | 403140050 |
| Coomassie Brilliant Blue G-250 | Thermofisher Scientific | 191480250 |
| Dithiothreitol (DTT) | Fisher Scientific | BP172 |
| Methanol | VWR | VWRC20903.550 |
| Ortho-Phosphoric Acid | Fisher Scientific | A260 |
| Pierce Protease Inhibitor Tablets | Thermofisher Scientific | A32955 |
| Precision Plus All Blue Ladder | Bio-Rad | 161037 |
| Sodium Chloride | Fisher Scientific | S671 |
| Sodium Dodecyl Sulfate | Fisher Scientific | BP166 |
| Sucrose | Fisher Scientific | BP 220-212 |
| TGX Stainfree 12% Gel | Bio-Rad | 1610185 |
| Tris (hydroxymethyl) Aminomethane | Fisher Scientific | BP152 |
| Tris(hydroxymethyl)aminomethane Hydrochloride | Fisher Scientific | BP153 |
| Triton-X 100 | Fisher Scientific | BP151 |
| Tween-20 | Fisher Scientific | BP337 |

**Table 2.** Western Blot Antibodies for Myelin Inclusionary and Exclusionary Markers.

| **Target** | **Acronym** | **Size (kDa)** | **Dilution Factor** | **Marker Type** | **Producer** | **Product Number** |
|---|---|---|---|---|---|---|
| ATP Synthase F1 Subunit | ATP5a | 60 | 2000 | Exclusionary | Abcam | ab140365 |
| β3-tubulin | TUBB3 | 50 | 2000 | Exclusionary | Abcam | ab78078 |
| β4-tubulin | TUBB4 | 50 | 2000 | Exclusionary | Abcam | ab11315 |
| Binding Immunoshock Protein | GRP78 | 78 | 2000 | Exclusionary | Abcam | ab140365 |
| Glial Fibrillary Acidic Protein | GFAP | 50 | 2000 | Exclusionary | DAKO | ZO334 |
| Glyceraldehyde-3-phosphate dehydrogenase | GAPDH | 36 | 2000 | Exclusionary | Abcam | ab8245 |
| Neurofilament Heavy Chain | NEFH | 180-220 | 2000 | Exclusionary | Abcam | ab8135 |
| Allograft Inflammatory Factor | IBA1 | 17 | 1000 | Exclusionary | Abcam | ab178846 |
| Sodium Potassium Pump | Na+/K+ ATPase | 100 | 2000 | Exclusionary | Abcam | ab140365 |
| Spectrin | SPTBN1 | 274 | 2000 | Exclusionary | Abcam | ab72239 |
| Myelin Associated Glycoprotein | MAG | 100 | 2000 | Myelin | CST | 9043S |
| Myelin Proteo-lipid Protein | PLP | 26, 20 | 2000 | Myelin | CST | 85971S |
| Myelin Oligodendrocyte Glycoprotein | MOG | 28 | 2000 | Myelin | CST | 96457S |
| Myelin Basic Protein | MBP | 21.5, 18.5, 17, 14 | 2000 | Myelin | Abcam | ab40390 |
| Sirtuin-2 | SIRT2 | 43 | 2000 | Myelin | Abcam | ab211033 |
| 2,3-cyclic nucleotide-3-phosphodiesterase | CNPase | 45, 48 | 2000 | Myelin | Abcam | ab227218 |
| Neurofascin-155 | NF155 | 140, 155 | 2000 | Myelin | Abcam | ab192399 |
| Anti-Mouse Starbright Blue 520 | - | - | 2500 | Anti-Primary | BIO-RAD | 12005867 |
| Anti-Rabbit Starbright Blu 700 | - | - | 2500 | Anti-Primary | BIO-RAD | 12004159 |

### Animals

All animal work was performed at the University of Northern British Columbia. Experimental procedures (#2020-16) were approved by the institutional Animal Care and Use Committee (ACUC) in accordance with the Canadian Council on Animal Care (CCAC). C57BL/J6 mice (RRID:IMSR_JAX:000664) were purchased from Jackson Laboratory (Bar Harbor, ME) to establish an in-house breeding colony. The animals were housed in groups of 2-4 per cage on a 12-hour light/dark cycle with ad libitum supply of food (Rodent Diet #5001; LabDiet Inc, St. Louis, MO) and water. Whole brain and isolated cortices (including cortex and corpus callosum) from adult mice were used for optimization of myelin fractionation. For aging analyses, myelin membranes were isolated from the cortices of 3, 12 and 20 month old mice (n = 4 per age group with 2 male and 2 female) corresponding to young, middle-age and aged phenotypes (41). Mice were anaesthetized in a closed chamber with 5% isoflurane in oxygen at a flow rate of 2L/min. A deep plane of anesthesia was confirmed by the absence of the pedal withdrawal reflex, and mice were subsequently euthanized by decapitation. Brain tissue was flash frozen in liquid nitrogen and stored at -80°C until further processing.

### Myelin Fractionation

Myelin was isolated using a discontinuous sucrose centrifugation protocol (Fig 1a), as previously described with minor modifications (42–44). All buffers contained 10 mM HEPES (pH 7.4), 10 mM dithiothreitol (DTT) & protease inhibitor cocktail (PI). Brain tissue was homogenized in 1M sucrose buffer using a Kimble 15 mL Dounce homogenizer (K885300-0015, DWK Life Sciences) using 7 strokes of the large clearance pestle and placed into a SW41 (14 x 89 mm) ultra-clear tube (#C14293). Tissue homogenate (TH) was subsequently layered with 0.85M and 0.35M sucrose. The gradient was centrifuged using a Beckman SW41 rotor (#333790) at 100,000 x g for 16 h at 4°C. Crude compact myelin (CC) and crude non-compact myelin (NC) were extracted from the 0.35/0.85M and 0.85/1M interface, respectively, then diluted ∼10 fold with HEPES buffer and centrifuged at 120,000 x g for 1 h at 4°C. The crude compact myelin underwent two rounds of osmotic shock by resuspension in HEPES buffer followed by centrifugation using a SW60 Ti rotor (#335649) at 120000 x g for 30 min at 4°C. A secondary discontinuous sucrose gradient was prepared by resuspending the crude compact myelin 1.0M sucrose buffer and subsequently layering with 0.70M, 0.62M and 0.35M sucrose. Myelin was floated up using a Beckman SW41 rotor (#333790) at 100000 x g for 4 h at 4°C. Compact myelin light (CL), medium (CM), and heavy (CH) were collected at the 0.35/0.62M, 0.62/0.7M, and 0.7/1.0M interfaces, respectively. The samples were diluted with HEPES buffer and washed as described above. CL and CM then underwent another two rounds of osmotic shock. For protocol validation, fractions were solubilized in SDS buffer (2% (w/v) SDS, 75 mM tris, 10 mM DTT and PI; pH 8.8) for characterization by western blot or in Triton-X 100 buffer (1% w/v TX100, 150 mM sodium chloride, 20 mM Tris; pH 7.5) for lipid assays. For protein analyses, myelin subfractions were lyophilized to determine dry mass by gravimetric analysis and then subsequently resuspended in high SDS buffer (8% (w/v) SDS, 150 mM NaCl, 25 mM Tris; pH 8.8). Samples were stored at -80°C until further analyses. For global lipidomics, a modified second gradient containing 0.35, 0.7, and 1.0M sucrose was used to combine the CL/CM fractions (0.35/0.7M interface) and pellets were stored under nitrogen gas at -80°C until lipid extraction.

### Western Blotting

Western blot was performed as previously described (44). Briefly, subfractions were then heated (95°C) with a modified Laemmli buffer (2% (w/v) SDS, 10% (v/v) glycerol, 0.01% (w/v) bromophenol blue, 1% β-mercaptoethanol, 62.5 mM Tris; pH 8.8). For initial characterization of myelin fractionation, 5 µg of protein was loaded onto 12% resolving gel (12% (w/v) 37.5:1 acrylamide: bis-acrylamide, 0.1% (w/v) SDS, 390 mM tris; pH 8.8) with a 5% stacking gel (5% (w/v) 37.5:1 acrylamide: bis-acrylamide, 0.1% (w/v) SDS, 130 mM tris; pH 8.8). Proteins were separated at 150V for stacking, followed 100V for resolving at 4°C. Gels were then equilibrated with a modified Towbin buffer (0.025% (w/v) SDS, 20% (v/v) methanol, 190 mM glycine, 25 mM tris; pH 8.8) before being transferred to 0.2 µm low fluorescence polyvinylidene fluoride membrane (PVDF; Amersham Hybond, #10600022; Cytiva) at 30 mA for 16 h at 4°C. The blot was blocked with 5% (w/v) skim milk in tris-buffered saline with Tween-20 (TBST; 0.1% (v/v) Tween-20, 140 mM NaCl, 20 mM Tris; pH 7.4) for 1 h at room temperature (RT). All primary antibodies were diluted in 5% skim milk in TBST and incubated overnight at 4°C. The blot was washed with TBST, incubated with the secondary antibody for 2 h at RT, and then washed again. Images were captured with a Chemidoc MP (Bio-Rad) using the Starbright 520 (475/30 ex, 532/28 em) or Starbright 700 (475/30 ex, 715/30 em) settings on automatic exposure.

For analysis of myelin protein over the age-span, western blots were performed as described above using 12% TGX Stain-Free gels (Bio-Rad; #1610185). Samples were loaded at 0.25-1 µg protein depending on the linear dynamic range of antibody detection (44). After electrophoresis, the gel was activated with UV for 45 seconds prior to transfer to PVDF. Blots were imaged using the stain-free filter (UV ex, 590/110 em) on manual exposure for total protein prior to probing for myelin antigens. Post-immunoprobing, fluorescence was captured on manual exposure setting with time based on lowest exposure time for each antigen. Image analysis was performed using Image Lab v6.1.0.07 (Bio-Rad). To quantify total protein, lane scan boundaries were set between the 15 kDa and 250 kDa molecular weight (MW) markers and background subtraction utilized a rolling disc (100 mm) such that a straight line was drawn from R_f_ = 0 to R_f_ = 1. For antigens, background subtraction utilized a disc size which spanned the entire signal (including doublet bands). The antigen fluorescence intensity was normalized to total protein, and fold change was expressed relative to the young age group. Statistical significance was determined by one-way ANOVA with Holm-Šidák post-hoc test (p < 0.05) using Sigmaplot v14.5 (Grafiti, Palo Alto).

### Lipid Assays

Total phospholipid (PL) and sterol (ST) content was measured using Phospholipid Assay Kit (Sigma; #MAK122) and Amplex Red Cholesterol Assay Kit (Invitrogen; #A12216), according to manufacturer’s protocols. Each protocol was scaled to a 384-well plate format, incubated at 37°C for 30 minutes and measured using a BioTek Synergy Neo2 plate reader (Agilent).

Concentrations were then interpolated using the respective standards and normalized to protein concentration. Statistical significance was determined by one-way ANOVA with Holm-Šidák post-hoc test (p < 0.05) using Sigmaplot v14.5.

### Global Lipidomics

Liquid chromatography-mass spectrometry (LC-MS/MS) was performed by The Metabolomics Innovation Centre (TMIC) of Canada (University of Alberta, Edmonton, AB, Canada) according to Buzatto et al (45, 46). Lipids were extracted using a modified Folch method. Prior to extraction, sample homogenate was spiked with NovaMT LipidRep Internal Standard Basic Mix containing 15 deuterated lipids (Nova Medical Testing, Inc., Edmonton, AB). The organic phase was collected and evaporated under nitrogen gas. Lipids were resuspended in NovaMT MixB (10 mM ammonium formate in 95:5 isopropanol/ water) and then diluted with NovaMT MixA (10 mM ammonium formate in 50:40:10 methanol/ acetonitrile/ water). A quality control (QC) sample made from pooling an aliquot from all experimental groups and biological replicates and processed in parallel.

Chromatographic separation and detection of analytes were performed using a Thermo Scientific Vanquish UHPLC linked to Bruker Impact II QTOF Mass Spectrometer using a Waters ACQUITY Premier CSH C18 column (100 x 2.1 mm, 1.7 μm). The mobile phase was composed of the NovaMT MixA and NovaMT MixB and analytes were resolved using the NovaMT 20-min-gradient (t = 0 min, 5% B; t = 10 min, 40% B; t = 18.8 min, 98% B; t = 20.5 min, 98% B) at a flow rate of 250 µL/min. Samples were injected at 4 and 12 µL for positive and negative mode, respectively. For mass spectrometry detection, the mass scan was set from 150-1500 m/z with a MS/MS collision energy ramping from 10 to 60 eV. Acquisition rate for MS was set to 1.44 Hz and 4-10 Hz for MS/MS.

Identification of putative lipids from MS/MS peaks was processed using NovaMT LipidScreener. The minimum intensity threshold was set to 3000 counts with a signal to noise ratio of 3; features that were not detectable for ≥ 50% of all sample injections were filtered out. The minimum peak length was 6 spectra with a retention time tolerance of 6 seconds for retention time correction and 4 seconds for alignment. The m/z tolerance was 20.0 ppm for peak picking, and 20.0 ppm and 5.0 mDa for alignment. A three-tiered identification approach was used based on nomenclature from the Lipid Maps database (47): Tier 1) MS/MS match score of ≥ 500, with precursor m/z error of ≤ 20.0 ppm and 5.0 mDa; Tier 2) MS/MS match score of ≤ 500, with precursor m/z error of ≤ 20.0 ppm and 5.0 mDa; and Tier 3) mass match with m/z error ≤ 20.0 ppm and 5.0 mDa. All identified features were first normalized to an internal standard with the nearest combination of lipid class similarity and then to the median intensity ratio found within each injection experiment. Tier 3 identifications lipids were then further filtered based on chain length (monoacyl >22 carbons; diacyl >44 carbons; triacyl > 54 carbons).

Multivariate and univariate statistical analysis was performed using MetaboAnalyst 6.0 (48). MS peaks that had the same molecular identification were combined as a singular intensity. For Principal Component Analysis, data was scaled by mean-centering and division by the standard deviation of each variable. Molecular identifications (fold change (FC) > 1.4 or < 0.7) were defined as significant at p < 0.05 with a false discovery rate (FDR) of q < 0.1 using one-way ANOVA with Fisher LSD post-hoc test. Lipid pathway analysis was performed using the web-based tool LipidSig 2.0 (49). Z-score was computed for each reaction set (with identified substrate and product) to determine pathway activation or suppression (p < 0.05; Z ≥ 1.645 or Z ≤ - 1.645). Plots comparing middle-age vs young, aged vs young, and aged vs middle-age were combined manually using Affinity Suite V2 (Serif Europe Ltd). Lipid ontology was performed using the web-based tool LION/web (50) using target-list mode, where the list of significantly altered lipids were compared to the universe list of all identified lipids (FDR q < 0.05).

## AUTHOR STATEMENTS

### Data availability

Data generated during this study are available from the corresponding author upon reasonable request.

### Author Contribution

KLF contributed to conceptualization, methodology, formal analysis, project administration and funding acquisition. VPL and CEJ contributed to methodology, investigation, formal analysis and visualization. SL contributed to investigation. KLF and VPL contributed to writing including original draft, review and editing.

## Acknowledgements

The authors would like to thank Dr. Liang Li and Dr. Shuang Zhao from The Metabolomics Innovation Centre (TMIC) for valuable consultation regarding methodology and data analysis. The authors also thank Sydney Cruch and Sianne Vautour for assistance with animal colony management.

