## Supplemental Figures S1-2 for "Lipidome of Compact Myelin from Cortical Regions of the Aging Murine Brain"

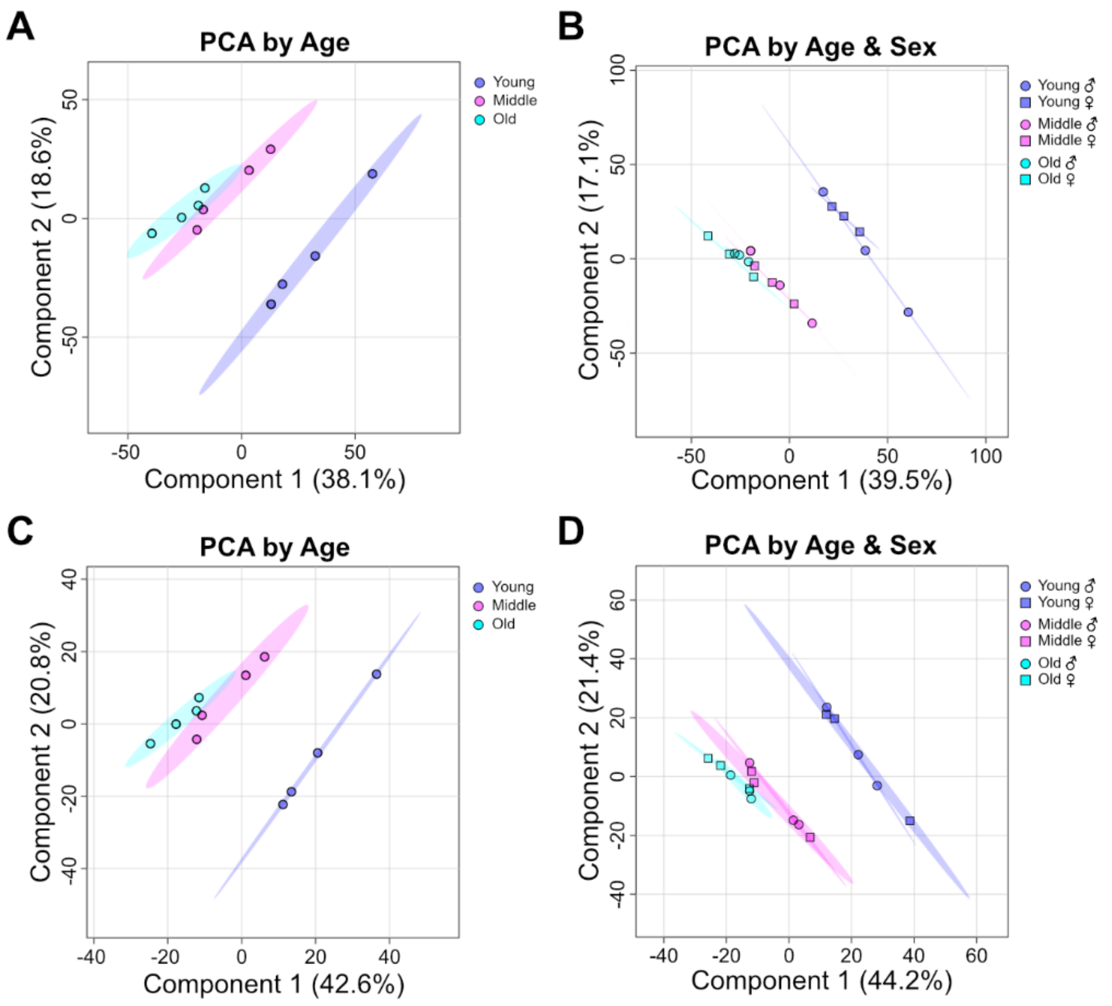


**Figure S1. PCA score plots of aging myelin lipidome.** PCA score plots displaying age-dependent (**A, C**) and age- and sex-dependent (**B, D**) clustering of unique molecular identifications from myelin lipidome in young (3 months; blue), middle-aged (12 months; magenta), and aged (20 months; cyan) mice (n = 4, n = 2/sex). A greater separation across age is observed that across sex; for preliminary analysis on sex differences, an average of the 2 biological replicates for each sex multiplied by 0.95 was used to obtain a third data point. Comparison of PCA score plots between dataset with inclusion of Tier 3 identifications (**A-B**; T1, T2, T3) versus exclusion of Tier 3 (**C-D**; T1, T2) shows minimal differences in clustering patterns.

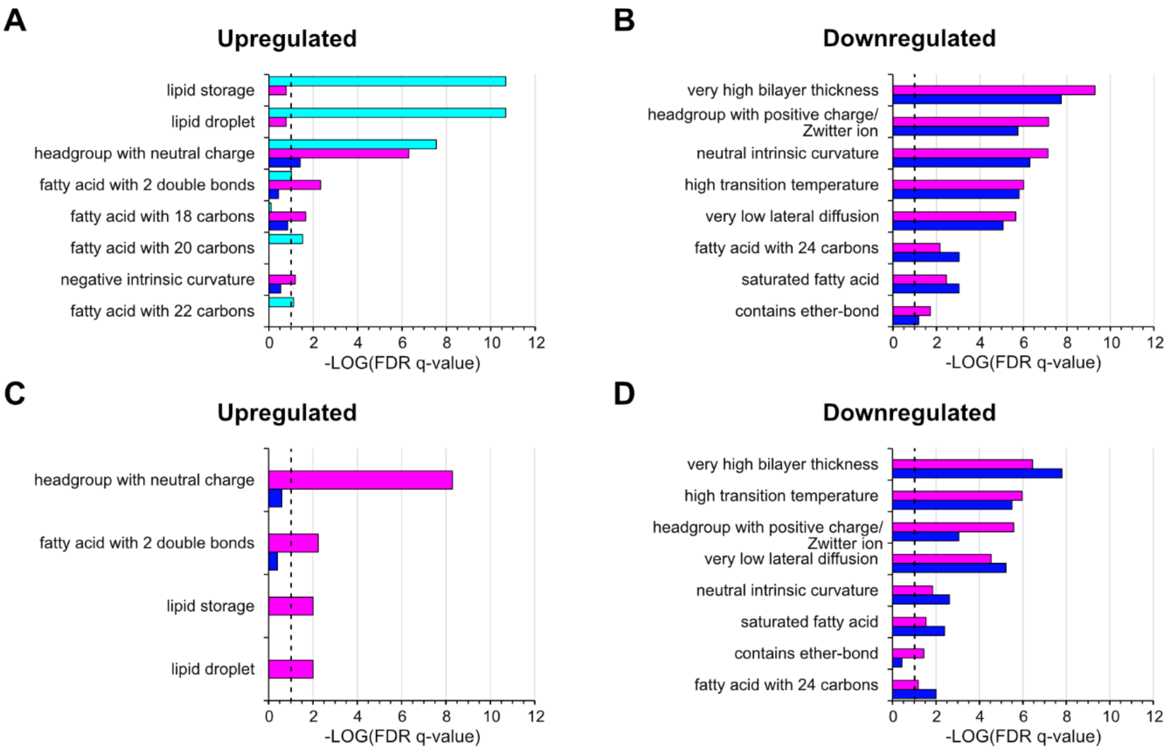


**Figure S2: Predicted alterations in lipid ontology terms in murine cortical myelin lipidome over aging.** Lipid ontology (LION) from LION/web analysis shows upregulated (**A, C**) and downregulated (**B, D**) terms for unique molecular identifications from myelin lipidome in young (3 months; blue), middle-aged (12 months; magenta), and aged (20 months; cyan) mice ($n = 4$, $n = 2$/sex; FDR $q < 0.1$). Comparison of PCA score plots between dataset with inclusion of Tier 3 identifications (**A-B**; T1, T2, T3) versus exclusion of Tier 3 (**C-D**; T1, T2) show similar outputs. Terms related to biochemical characteristics of lipids, membrane biophysical characteristics of membrane and lipid trafficking are shown (all terms including those that refer to specific lipid species or subclasses are reported in Table S3).
